# Identification and Removal of Doublets with DoubletDecon

**DOI:** 10.1101/2020.04.23.058156

**Authors:** Erica A. K. DePasquale, Daniel Schnell, Kashish Chetal, Nathan Salomonisi

## Abstract

Retention of multiplet captures in single-cell RNA-sequencing (scRNA-seq) data can hinder identification of discrete or transitional cell populations and associated marker genes. To overcome this challenge, we created DoubletDecon to identify and remove doublets, multiplets of two cells, by using a combination of deconvolution to identify putative doublets and analyses of unique gene expression. Here we provide the protocol for running DoubletDecon on scRNA-seq data.

For complete details on the use of this protocol, please see DePasquale et al. (2019) (https://doi.org/10.1016/j.celrep.2019.09.082).

**GRAPHICAL ABSTRACT:** 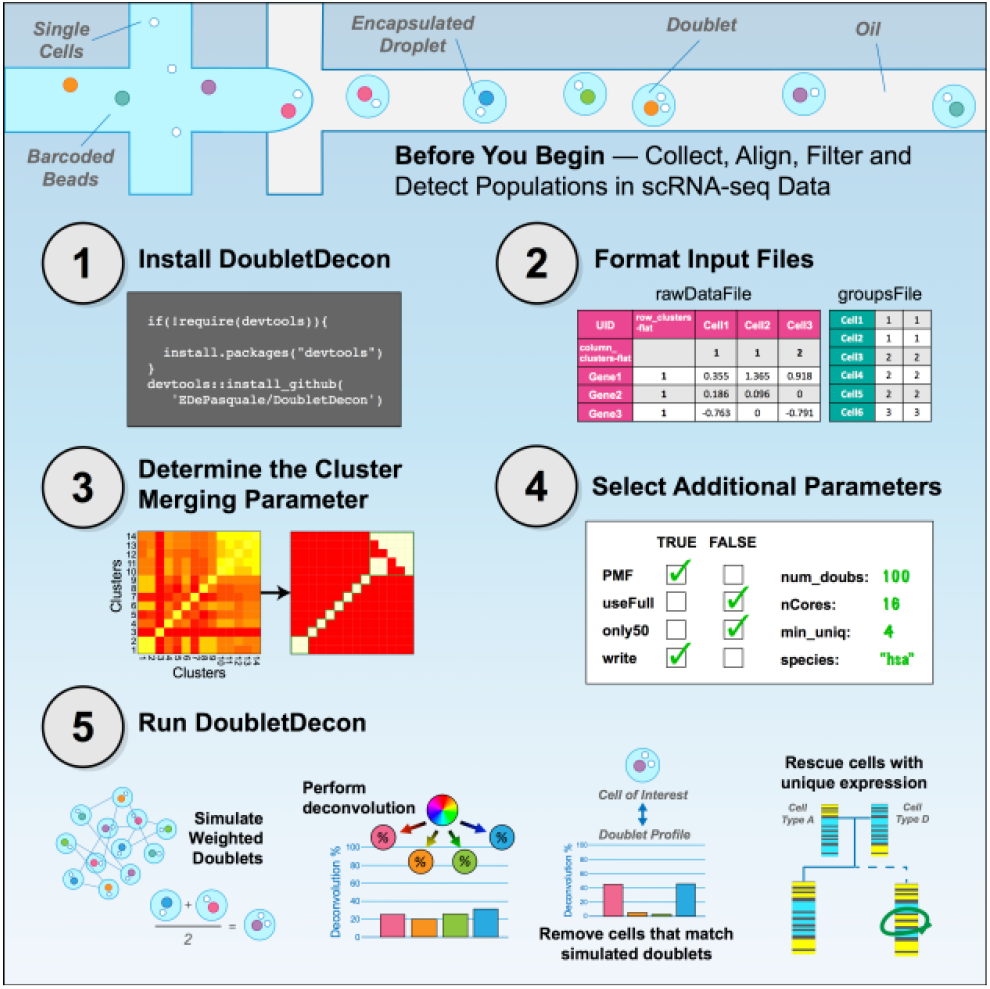

## BEFORE YOU BEGIN

### A. Data Collection

DoubletDecon was designed to detect doublets in droplet-based scRNA-seq data, such as those produced using the 10X Genomics and Drop-Seq platforms and emerging technologies (DePasquale et al., 2019). The software assumes that doublet cell populations do not possess unique gene expression relative to other cell clusters. Due to DoubletDecon’s ability to rescue cells based on unique gene expression relative to predicted doublet contributors, we recommend sequencing cells to a minimum depth of 1,000 unique molecular indexes (UMIs) or 50,000 non-unique reads per cell.

### B. Read Alignment

We recommend aligning your reads according to the manufacturer’s recommendations for all single-cell sequencing technologies.

### C. Filtering

As with the read alignment suggestions, we recommend filtering genes based on the manufacturer’s recommendations. It is important to note that the results from all doublet detection algorithms, including DoubletDecon, may differ dramatically based on the level of gene filtering (DePasquale et al., 2019). Although we recommend at least 500 genes expressed per cell in most cases, DoubletDecon has been shown to perform well with as low as 150 genes per cell barcode.

### D. Population Detection

DoubletDecon has been described as a “cell-state” or “cell-type aware” application, meaning that the workflow is applied after obtaining cell populations from an unsupervised or supervised analysis. This allows for the software to quickly identify overlapping cell populations, select cells for synthetic doublet creation, and identify marker genes from these clusters. DoubletDecon is compatible with results from diverse supervised and unsupervised population detection tools, including the software Seurat (Satija et al., 2015) and Iterative Clustering and Guide Gene Selection (ICGS) (Olsson et al., 2016; Venkatasubramanian et al., 2019). The output of both applications is directly supported in DoubletDecon, though alternative clustering methods may be used as long as the files are formatted appropriately. The following variables should be considered when running population detection algorithms for the purposes of doublet detection:

1. <u>Cluster Number</u> – Both ICGS and Seurat can automatically determine the optimal number of transcriptionally distinct clusters in a dataset, with both tools allowing the user to manually adjust cluster numbers using resolution parameters. ICGS version 2 should not identify full doublet cell clusters, which would be problematic for DoubletDecon, because it requires each cell population to have unique marker genes. Although DoubletDecon includes a cluster merging step to mitigate the effects of overly granular clustering, overly broad clustering should be corrected prior to performing doublet removal. Care should be taken to correct this by increasing the resolution before using DoubletDecon. For more information of what clustering should look like for use with DoubletDecon, please see the description in Step 3 of the protocol below.
2. <u>Cell Cycle</u> – DoubletDecon has an optional parameter to remove cell cycle-enriched gene clusters prior to initial doublet predictions (removeCC = TRUE), but it may be beneficial to negate the effects of cell cycle stage on initial cell clustering and population prediction. These genes can cause separation of highly related cell types in the absence of differential gene expression outside of cell cycle genes. As such, removing cell cycle effects at this stage may improve doublet detection.

### E. Dataset Merging

When using data integration techniques, we recommend running any doublet removal tool, including DoubletDecon, on each sample prior to integration.

## MATERIALS AND EQUIPMENT

- Data (single-cell RNA-sequencing normalized reads and clustering – see ‘Population Detection’ in ‘Before You Begin’).
- R software and required packages. The required packages will be automatically installed the first time you run DoubletDecon. While older versions of the R software and associated packages may work correctly with DoubletDecon, the authors use R v3.5.0 and the following packages at the indicated versions when writing this protocol:

- DeconRNASeq (v1.26.0)
- gplots (v3.0.1.1)
- dplyr (v0.8.3)
- MCL (v1.0)
- clusterProfiler (v3.12.0)
- mygene (v1.20.0)
- tidyr (v1.0.0)
- R.utils (v2.9.0)
- foreach (v1.4.7)
- doParallel (v1.0.15)
- stringr (v1.4.0)
- plyr (v1.8.4)
- R Studio software and the following packages for use with the DoubletDecon UI (MacOS only):

- shiny (v1.3.2)
- shinyDirectoryInput (v0.2.0)
- shinyFiles (v0.7.3)
- ggplot2 (v3.2.1)
- reshape2 (v1.4.3)
- shinyjs (v1.0)
- V8 (v2.3)
- umap (v0.2.3.1)
- shinycssloaders (v0.2.0)
- scales (v1.0.0)
- png (v0.1-7)
- Hardware

- Local – Memory: 8GB required, 16GB recommended; Processors: 1 required, 4 recommended.
- Computational Cluster – Memory: >64GB recommended; Processors: >8 recommended for parallel processing and large datasets.

**Pause Point:** If you want to pause at any time during this protocol, please save your work and R session using the following command within the R or R Studio console:

~~~
          >save.image(“Your Experiment Name Here.Rdata”)
~~~

The session can be reloaded by using the following command:

~~~
          >load(“Your Experiment Name Here.Rdata”)
~~~

## STEP-BY-STEP METHOD DETAILS

### Step 1. Installing DoubletDecon

#### Timing: 5 minutes

Full installation of DoubletDecon includes downloading the DoubletDecon package from GitHub as well as the access files for the user interface.

1. Install DoubletDecon by running the following code:

~~~
>if(!require(devtools)){
>        install.packages(“devtools”) # If not already installed
>}
>devtools::install github(‘EDePasquale/DoubletDecon’)
~~~
2. To access the user interface:

a. Download the DoubletDecon GitHub repository at https://github.com/EDePasquale/DoubletDecon/archive/master.zip.
b. Unzip the directory
c. Inside the directory, click on the ‘DoubletDecon.app’ to start the application. This will automatically start R and load the user interface in a browser window.

### Step 2. Formatting input files for use with DoubletDecon

#### Timing: 5–30 minutes

DoubletDecon accepts input from both ICGS and Seurat unsupervised clustering pipelines either directly (ICGS) or with use of a conversion function (Seurat).

**Note**: Example files for testing DoubletDecon can be found at the project GitHub repository in the folder “Seurat_Conversion_Files”. This folder contains the Seurat object that has been generated following the below instructions (Variant 2) as well as the three output files that have been converted to ICGS version 2 format using the DoubletDecon function Improved_Seurat_Pre_Process().

#### Variant 1. Iterative Clustering and Guide Gene Selection (ICGS) version 2

1. DoubletDecon is designed to natively work with ICGS files, so no reformatting is necessary. The files that are needed for running DoubletDecon can be found navigating through the paths listed below:

a. ICGS marker gene expression file: ~/Project_Name/ICGS-NMF/FinalMarkerHeatmap_all.txt
b. ICGS groups file: ~/Project_Name/ICGS-NMF/FinalGroups.txt
c. (Optional) Full expression file: ~/Project_Name/ExpressionInput/exp.Project_Name.txt

#### Variant 2. Seurat version 3

1. Seurat files can be converted into ICGS-like file formats using the Improved_Seurat_Pre_Processing() function using the following code:

~~~
>newFiles=Improved Seurat Pre Process(seuratObject, num genes=50,
  write files=FALSE
~~~ The required parameters are:

a. <u>seuratObject</u>: Seurat object following a protocol such as https:ZZsatijalab.org/seurat/v3.1Zpbmc3k_tutorial.html or using the ‘seurat-3.0.R’ script provided in the DoubletDecon GitHub repository.
b. <u>num_genes</u>: Number of genes for the top_n function. Default is 50. This parameter determines the number of marker genes that are output for each cell cluster. Fewer than 50 marker genes per cluster is likely to reduce the performance of DoubletDecon.
c. <u>write_files</u>: Save the output files to .txt format. Default is FALSE. It is recommended to set this to TRUE when the size of the dataset is larger or when you want to rerun DoubletDecon more than once on the same dataset, as the Improved_Seurat_Pre_Processing() function can take some time.
2. The outputs of this function are three files directly equivalent in format to the ICGS files described above:

a. ICGS marker gene expression file: newFiles$newExpressionFile
b. ICGS groups file: newFiles$newGroupsFile
c. (Optional) Full expression file: newFiles$newFullExpressionFile

**Note**: This is a good time to compare your input files to example files provided in the DoubletDecon GitHub repository (https://github.com/EDePasquale/DoubletDecon) to ensure that the conversion completed correctly. This is particularly important as unsupervised clustering software packages are often updated, leading to format differences from what was initially anticipated.

### Step 3. Determining the cluster merging parameter (ρ’)

#### Timing: ~15 minutes

Cluster merging is a unique feature of DoubletDecon that allows the user to overcome overly-granular clustering of cells in the pre-processing steps to establish transcriptionally distinct clusters for use with DoubletDecon. The algorithm accomplishes this by calculating a Pearson correlation threshold for the similarity of clusters determined by the user input value of ρ’ (“rho prime”). Two methods can be used to identify an appropriate ρ’ value: manually or through the user interface. Further details and considerations for this step are presented in the STAR Methods of the original DoubletDecon paper (DePasquale et al., 2019).

#### Variant 1. Manual Identification

1. Begin by running DoubletDecon with default parameters, including a ρ’ value of 1. This ρ’ value results in the cluster similarity threshold being set to the cluster correlation mean plus 1 standard deviation, and is generally a good starting point.
  a. The code for this is:

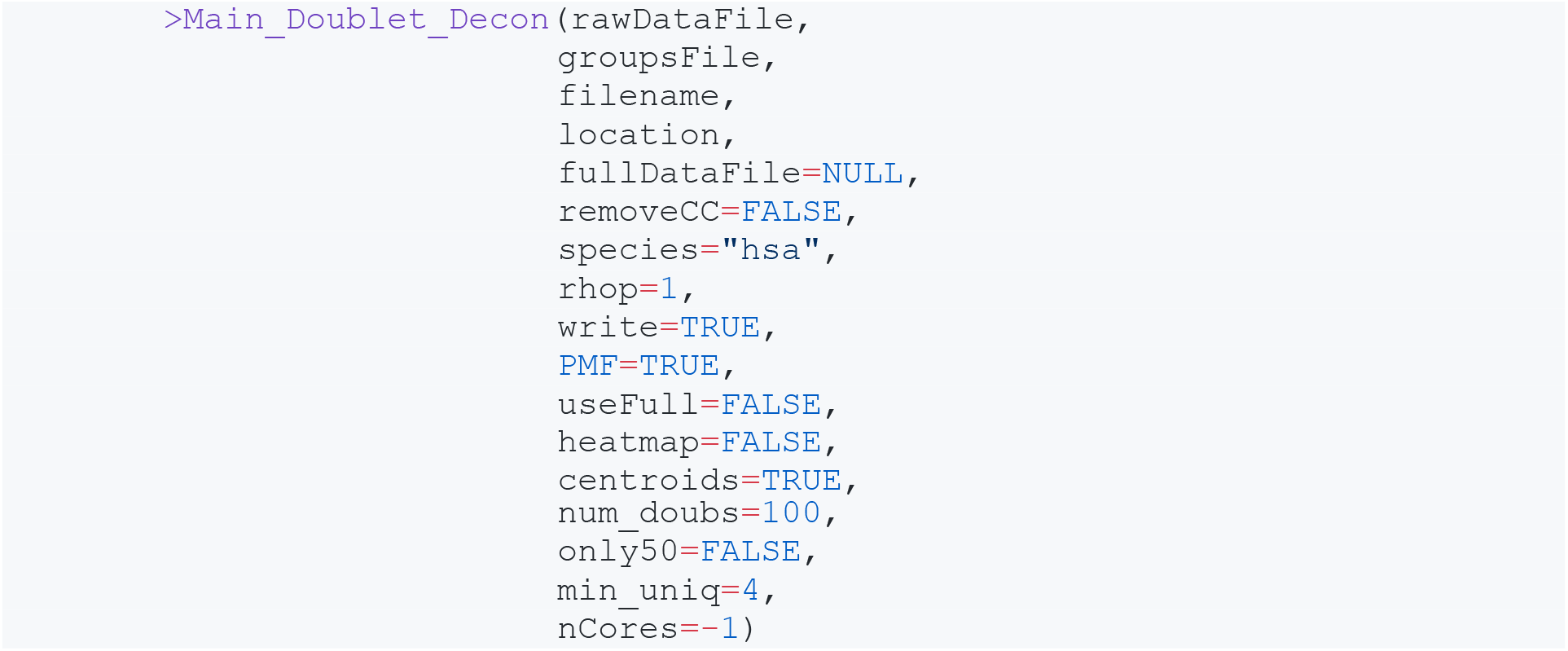
2. When DoubletDecon completes the cluster merging step (denoted by printing of “Creating synthetic doublet profiles…” in the console and log file), terminate DoubletDecon.
3. Examine the clustering merging heatmap that was automatically generated in conjunction with previously identified cluster labels to assess the appropriateness of merging. The example below in Figure 1 shows the result of various ρ’ values in the mouse hematopoietic progenitor dataset from Figure 3 of the original DoubletDecon paper (DePasquale et al., 2019). **Figure 1A** shows no cluster merging, which may cause a loss of sensitivity in DoubletDecon due to the biological similarity of the HSCP-1 and HSCP-2 clusters as well as the similarity of the monocyte-dendritic cell precursor (MDP) and monocyte progenitor (Mono) clusters. **Figure 1B** shows appropriate merging of these clusters. Finally, **Figure 1C** shows over-merging of clusters, with the hematopoietic stem cell progenitors (HSCP) being combined with the megakaryocyte progenitors (Meg) and the monocyte progenitors combined with the granulocyte progenitors (Gran).
4. When merging clusters, the goal is to combine highly similar clusters to improve the formation of reference clusters for deconvolution and to generate transcriptionally distinct synthetic doublets, both of which improve the overall accuracy of the method. However, by merging two clusters the software will no longer be able to detect doublets formed between those two clusters. This is why a thorough understanding of the cell cluster identifications and relevant biology are critical to using DoubletDecon, as it may be very important to remove doublets of some classes while retention of other doublet types may not affect downstream analyses.
5. Repeat the above process while adjusting ρ’ down (if too few clusters are merged) or up (if too many clusters are merged) by 0.1 increments until an acceptable ρ’ value is reached

**Figure 1.**
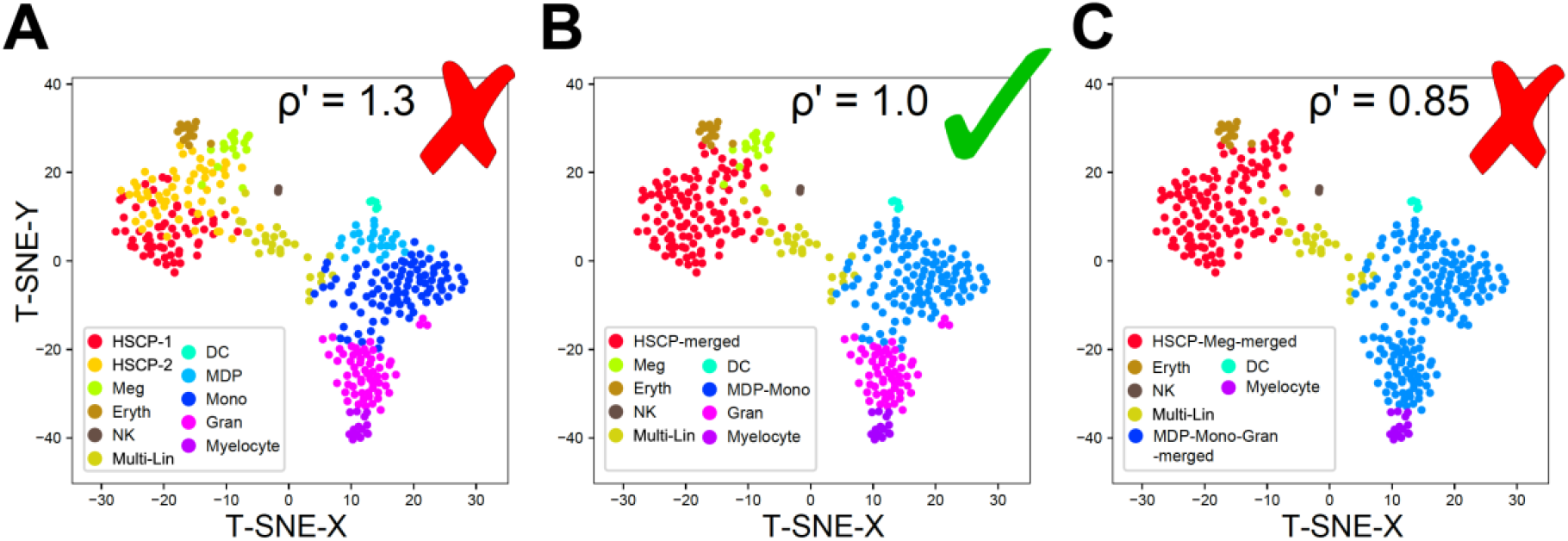
Cluster Merging Parameter Selection Examples. Mouse hematopoietic progenitor cells sequenced on the Fluidigm C1 platform with clusters identified by ICGS version 1 and cluster merging performed at various ρ’ values. (A) Too little merging (ρ’=1.3), (B) Appropriate merging (ρ’=1.0), and (C) Too much merging (ρ’=0.85).

#### Variant 2. Interface Identification

1. Begin by starting the DoubletDecon desktop application and uploading all relevant data files. Keep the remaining parameters at their default values.
2. Select the “Test for rho-prime values” button at the bottom of the initial screen. A new tab will automatically open when processing is completed, which will display a number of cluster merging heatmaps along with the associated ρ’ values used to generate those heatmaps. **Note:** Please be patient, as the processing for this step may take several minutes.
3. Complete Variant 1, steps 3 and 4 to decide on the appropriate ρ’ value for your experiment. Please note this value for later use.
4. Close and restart the DoubletDecon application before moving on to the following steps.

**Critical: The results of DoubletDecon are highly dependent on the selection of an appropriate ρ’ value! Please ensure that you understand the cell types/states of the clusters you are merging prior to selecting which clusters to merge.**

### Step 4. Selecting additional parameters

#### Timing: ~5 minutes

While the cluster merging parameter chosen in Step 2 of this protocol is the most critical for accurate and reliable performance of DoubletDecon, other parameters may affect the results and their values should be carefully considered. Listed below are the remaining parameters to choose, listed in descending order of importance in most test cases, with default values and tips for selection provided.

1. <u>only50</u> – this parameter, set to FALSE by default, determines whether the synthetic doublets are created with only a 50%/50% average of two parent cells (only50 set to TRUE) or if 70%/30% and 30%/70% weighted averages are included as well (only50 set to FALSE). In most cases, ‘only50’ set to FALSE increases the sensitivity of DoubletDecon to detect doublets at a minor decrease of specificity. However, datasets that are known to contain a large proportion of cells from transitional populations or those with more continuous cell definitions will likely have a higher accuracy when ‘only50’ is set to TRUE, assuming a small decrease in sensitivity.
2. <u>PMF</u> – this parameter, set to TRUE by default, ‘rescues’ cell clusters from inappropriate removal by checking for unique gene expression that cannot be explained as a combination of any of the remaining cluster expression profiles. The name is an abbreviation of “pseudo MarkerFinder”, as MarkerFinder is a function in AltAnalyze that performs a similar unique gene expression check (Olsson et al., 2016; Venkatasubramanian et al., 2019). In nearly all cases, rescue of erroneous identified clusters improves the accuracy of doublet detection, though the results of this ‘rescue’ should be reviewed for rescue of non-biologically feasible doublets.
3. <u>useFull</u> – this parameter, set to FALSE by default, checks for unique expression during the rescue set of DoubletDecon (when ‘PMF’ is TRUE) in all of the genes of the expression file. By leaving this parameter as FALSE, DoubletDecon only uses those genes included in the ‘rawDataFile’ input, which should be a set of genes selected to best define each cell cluster. When using useFull as TRUE, special care will need to be taken on selecting the appropriate value of ‘min_uniq’ (see below). In most cases, the use of the full gene set is unnecessary, as most cluster-defining genes are included in the ‘rawDataFile’ input, though unique gene expression may be found in one of the genes not represented. As such, we recommend testing your data with the ‘useFull’ option set to TRUE and an appropriate ‘min_uniq’ value selected if suspected progenitor or transitional clusters are removed.
4. <u>centroids</u> – this parameter, set to FALSE by default, is the parameter for determining whether cluster centroids or medoids are used to define the references for the deconvolution ‘remove’ step of DoubletDecon. Medoids (‘centroids’ set to FALSE) should be used when some cell clusters contain a large (>40%) proportion of suspected doublets when centroids may skew the result. However, centroids (‘centroids’ set to TRUE) better represent the data when this case is not met and is the recommended choice in most cases.
5. <u>num_doubs</u> – this parameter, set to 100 by default, is the number of synthetic doublets generated for each pair of clusters during the synthetic doublet generation step of DoubletDecon. This value, along with 250 and 500, gave similar and consistent results when testing the Demuxlet and Cell Hashing datasets in the paper, though a larger value should be chosen in very large datasets. A good rule of thumb would be approximately 10% of the number of cells in the dataset. Note: increasing this number greatly increases the run time of DoubletDecon, especially in combination with only50 set to FALSE (see below).
6. <u>removeCC</u> – this parameter, set to FALSE by default, checks for gene clusters enriched in cell cycle genes via KEGG analysis. When setting this parameter to TRUE to remove cell cycle gene clusters, the user will also be required to input the appropriate ‘species’ (see below) and have an internet connection. The presence of cell cycle clusters may confound the identification of doublets due to shared genes expressed between clusters. In most cases, cell cycle removal only improves doublet detection, though in some cases the cell cycle genes are what define a cluster (e.g. studying cell metabolism in tumors) and should be retained.
7. <u>nCores</u> – this parameter, set to −1 by default, is used during the ‘rescue’ step of DoubletDecon and indicates how many cores should be used when parallel processing of the full gene list is selected. The default value of −1 triggers automatic detection of core and will work for most purposes. When using a computational cluster, setting this value to the number of available cores designated in a particular job will eliminate related errors.
8. <u>min_uniq</u> – this parameter, set to 4 by default, indicates the minimum number of genes that must be uniquely expressed per potential doublet cluster to be ‘rescued’ during the final step of DoubletDecon. When using the full gene list (when ‘useFull’ is TRUE), the ‘min_uniq’ value should be set to 30, which was guided by the verified mouse non-doublet evaluation dataset in the original DoubletDecon publication (see STAR Methods for more details) (DePasquale et al., 2019). These values were chosen to correct for multiple tests and reduce the risk of false positive genes labeled as uniquely expressed. Increasing this number decreases the number of putative doublet clusters that are rescued.
9. <u>species</u> – this parameter, set to “hsa” by default, is used only when cell cycle genes are removed (when ‘removeCC’ is TRUE). The correct species should be indicated by the official three letter code to allow for accurate conversion of gene identifiers to use by the Enrichr module.
10. <u>write</u> – this parameter, set to TRUE by default, gives permission for all text-based output files of DoubletDecon to be written to the hard drive. DoubletDecon will over-write previous files saved with the same name, so please use a unique ‘filename’ name for each run. These files include intermediate results files from the primary steps of DoubletDecon, final doublet and non-doublet group files, and doublet and non-doublet expression files. Setting this parameter to FALSE should only be used when testing large number of parameter combinations while using an alternative method of storing the results.
11. <u>heatmap</u> – this parameter, set to TRUE by default, causes DoubletDecon to generate expression heatmaps for the original data, the final non-doublets, and the final doublets. This is useful for visually evaluating the DoubletDecon-predicted doublets and non-doublets overall accuracy when obvious doublet expression patterns are visible in the data. In extremely large datasets, this parameter should be set to FALSE during testing and heatmaps should only be generated when the final parameters are selected to minimize computational burden.

### Step 5. Running DoubletDecon

#### Timing: 5-30 minutes

After selecting all of the initial parameters and generating the formatted input files, the final step is to run DoubletDecon. DoubletDecon can be used either in the R console (or as part of a larger script) or with the user interface of the desktop application. Both methods produce the same doublet predictions, but the desktop application automatically generates interactive UMAPs and bar charts for visualization of the results that are not available using the command line version. Additional details for running DoubletDecon and updates to the parameters can be found at the project GitHub repository (https://github.com/EDePasquale/DoubletDecon).

#### Variant 1. Console

1. Load DoubletDecon by calling:

~~~
      >library(DoubletDecon)
~~~
2. Set your working directory (using setwd()) to the DoubletDecon folder containing the code used to run DoubletDecon. This is necessary for accessing the conversion files for cell cycle gene removal.
3. Run DoubletDecon by calling the Main_Doublet_Decon() function with all of the above parameters clearly defined as shown in the example below:

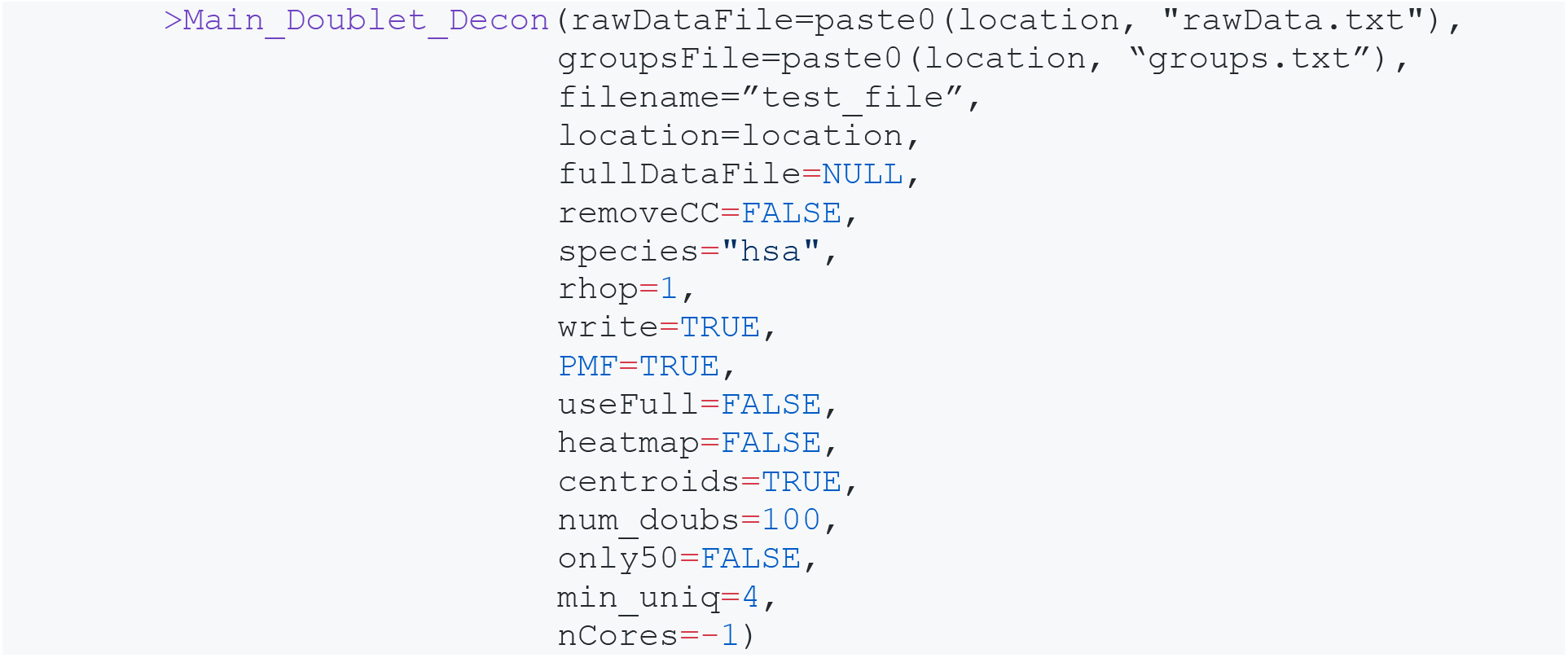

a. When using Seurat files, the ‘rawDataFile’, ‘groupsFile’, and (optional) ‘fullDataFile’ should be provided as:

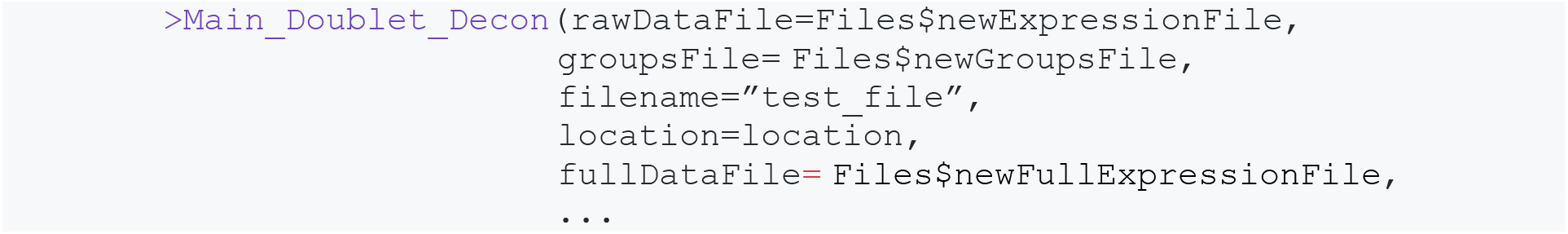

where ‘Files’ is the resulting object from the Improved_Seurat_Pre_Process() function.

#### Variant 2. Desktop Application

1. Begin by starting the DoubletDecon desktop application.
2. Choose a project name (or use the one automatically generated), which will serve as your ‘filename’ parameter, and select the directory containing your input files (the ‘location’ parameter).
3. Choose whether you wish to use ICGS files or Seurat files and upload appropriately.

a. When using Seurat, Option 1 is preferred and is the Improved_Seurat_Pre_Process() function, which takes as input a Seurat object prepared as described in Step 1 of this guide. Option 2 is legacy option that we have retained for people who have already generated the three necessary input files separately.
4. Input the remaining parameters decided in Step 3, keeping in mind that some of the parameters are not visible until necessary conditions are met (i.e. the dialog for uploading the full expression file will not be available until ‘Use full gene list?’ is set to TRUE.
5. Select the ‘Run DoubletDecon’ button at the bottom of the screen, which will run DoubletDecon with the given parameters. **Note**: The ‘Generate function call only’ button at the bottom of the user interface takes the provided parameters and generates code for running the same set of parameters in the console. The is especially useful if you wish to run DoubletDecon multiple times in a loop (see Optional Step 6) to improve accuracy.

**Pause Point:** Once you know the parameters you wish to use and have successfully run DoubletDecon, you may find this to be a good place to pause and evaluate the results before proceeding with the optional steps.

### Optional Step 6: Running DoubletDecon Multiple Times

#### Timing: 5–30 minutes

Due to the random nature of synthetic doublet generation in DoubletDecon, results will vary slightly across runs. To ensure only high-confidence doublets are predicted and subsequently removed from the input data, DoubletDecon should be run multiple times with the same or slightly varying settings. Following this, the intersection of the DoubletDecon doublet predictions could be used for further downstream analysis.

1. Run DoubletDecon and evaluate the results as described in the following section on Expected Outcomes.
2. Store the list of predicted doublets in a text file or R object. This list will be the row names of the “Final_doublets_groups” output of DoubletDecon.
3. Repeat steps 1 and 2 the desired number of times (recommended 20) and record the predicted doublet results from each run.
4. Identify the intersection of predicted doublets by loading the file or object into R as a list object. Using the intersect() function in R with each list of cell names as parameters in the function call will result in an intersection of the cell names provided (for more details on how to use the intersect() function, see https://stat.ethz.ch/R-manual/R-devel/library/base/html/sets.html).

### Optional Step 7: Assessing Consensus with Other Doublet Detection Methods

#### Timing: ~60 minutes

While DoubletDecon has been shown to perform comparably to alternative computational doublet detection approaches, using a combination of algorithms allows the user to prioritize sensitivity or specificity depending on the research question. Two options for this additional step would be to run two or more doublet detection algorithms and use the union or intersection of the predictions for downstream analyses. This process is very similar to what is described in Optional Step 6.

#### Variant 1. Intersection of doublet predictions

1. Run DoubletDecon and evaluate the results as described in the following section on Expected Outcomes.
2. Store the list of predicted doublets in a text file, Excel spreadsheet, or R object. This list will be the row names of the “Final_doublets_groups” output of DoubletDecon.
3. Run alternative computational doublet detection methods according to the developer’s instructions and save the lists of predicted doublets.
4. Identify the intersection of predicted doublets with one of the following methods:

a. If using an Excel spreadsheet to store predicted doublets, copy these cells into an interactive Venn Diagram generator, such as Venny (https://bioinfogp.cnb.csic.es/tools/venny/), taking care to place each algorithm’s predicted cells as separate lists. Click on the center of the Venn Diagram to generate a list of only cells predicted in all doublet detection algorithms.
b. If using a text file or R object, load the file or object into R as list objects. Using the intersect() function in R with each list of cell names as parameters in the function call will result in an intersection of the cell names provided (for more details on how to use the intersect() function, see https://stat.ethz.ch/R-manual/R-devel/library/base/html/sets.html).

#### Variant 2. Union of doublet predictions

1. Run DoubletDecon and evaluate the results as described in the following section on Expected Outcomes.
2. Store the list of predicted doublets in a text file, Excel spreadsheet, or R object. This list will be the row names of the “Final_doublets_groups” output of DoubletDecon.
3. Run alternative computational doublet detection methods according to the developer’s instructions and save the lists of predicted doublets.
4. Identify the union of predicted doublets with one of the following methods:

a. If using an Excel spreadsheet to store predicted doublets, simply paste together the lists of predicted doublet cell names to create the union of lists.
b. If using a text file or R object, load the file or object into R as list objects. Using the union() function in R with each list of cell names as parameters in the function call will result in a union of the cell names provided (for more details on how to use the union() function, see https://stat.ethz.ch/R-manual/R-devel/library/base/html/sets.html).

**Note**: Cell names containing special characters, such as “.”, “_”, “-”, “/”, etc. may have been altered during the DoubletDecon or alternative algorithm processes. Please ensure that all lists of cells have the same naming conventions following doublet prediction before combining results.

## EXPECTED OUTCOMES

### Number of Predicted Doublets

DoubletDecon is designed to predict doublets based on gene expression signatures alone and does not rely on statistical estimation of doublet count as an explicit parameter. However, it would be prudent to review the results of DoubletDecon for extreme over-estimation of doublet count.

Figure 2 shows UMAP projections for an example single-cell RNA-sequencing experiment in fewer than 30% of captures are expected to be doublets based on loading during sequencing. DoubletDecon predicted doublets are colored blue while predicted non-doublets are colored gray. When faced with the high proportion of predicted doublets shown in **Figure 2A**, the investigator can choose to re-run DoubletDecon with more restricted parameters to limit the amount of false positives predicted by the software. After modifying the ‘min_uniq’ and ‘only50’ parameters, the resultant doublet prediction numbers were more in line with statistical estimates and the predicted doublets were more frequently found at the intersection of two unrelated clusters, which resulted in an improved accuracy (**Figure 2B**). The parameters outlined in Step 4 of this protocol are listed in descending order of importance to the final doublet predictions. As such, parameters at the top of the list should be adjusted first to refine predictions. Additionally, the investigators could choose to perform Optional Step 7 with two doublet detection methods and use only the intersection of doublet predictions for their final data.

**Figure 2.**
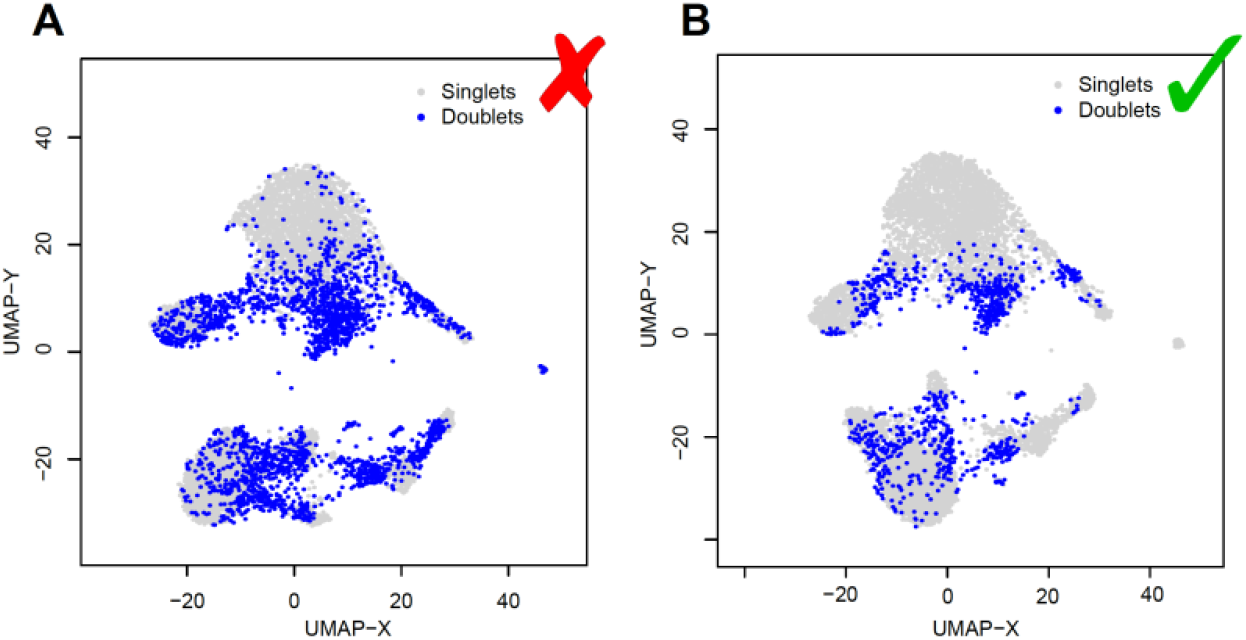
Number of Predicted Doublets Output Example. *In silico* identified doublet cell profiles obtained from the Dexmulet software, which identifies cells with a combination of genomic variants associated with the 8 profiled single-cell donors to find cellular barcodes with hybrid genotype profiles. (A) An example UMAP showing excessive doublet predictions with the initially provided DoubletDecon parameters, (B) An example UMAP showing refined doublet predictions following modification of parameters.

### Output Files and Formatting

The outputs of DoubletDecon are a series of .txt files containing information on the intermediate and final doublet predictions of the program when the ‘write’ parameter is set to TRUE. The following descriptions of output files have ‘TEST’ given as the filename while the names of your files will differ depending on your ‘filename’ parameter input. In addition to the following .txt files, users can save heatmaps, plots, and charts summarizing the results in the DoubletDecon UI.

#### Intermediate Results

1. <u>“data_processed_TEST.txt”</u> – This file (and associated data frame) is the initial normalized expression matrix that has been processed to include cluster identification rows and columns, renumber clusters in ascending numerical order, replace special characters such as “.”, “_”, “-”, or “/”, and optionally remove cell cycle-enriched gene clusters.
2. <u>“groups_processed_TEST.txt”</u> – This file contains cell-cluster associations following the Clean_Up_Input() step of DoubletDecon.
3. <u>“Synth_doublet_info_TEST.txt”</u> – This file contains information in which two cells were combined for each synthetic doublet generated in the Synthetic_Doublets() function. The first two columns are the names of the chosen cells and the second two columns are their cluster names, which may include cluster merging.
4. <u>“DRS_doublet_table_TEST.txt”</u> – This file contains the results of the initial ‘Remove’ step in DoubletDecon following the “Is_A_Doublet” function. While this file may be important for understanding the decision making behind DoubletDecon’s initial predictions, this is NOT the final doublet prediction list from DoubletDecon. The description of each column is as follows: name of cell, correlation to most similar cluster, most similar cluster number or name, logical value indicating whether this cell is initially considered a doublet, and the original cluster number or name for the cell.
5. <u>“DRS_results_TEST.txt”</u> – This file contains the results of the deconvolution step in DoubletDecon, with rows representing cells and columns representing clusters. The values in this file are predictions of the percent make up of each cell, with each row summing to 100%.
6. <u>“data_processed_reclust_TEST.txt”</u> – This expression file is very similar to the “data_processed_TEST.txt” file, but cells (columns) have been regrouped according to the initial doublet predictions in the ‘Remove’ step.
7. <u>“groups_processed_reclust_TEST.txt”</u> – This file is very similar to the “groups_processed_TEST.txt” file, but cells (rows) have been regrouped according to the initial doublet predictions in the ‘Remove’ step.
8. <u>“new_PMF_results_TEST.txt”</u> – This file contains information for the final doublet predictions from DoubletDecon following the ‘Rescue’ step, though the file itself does not contain novel information and is only used under certain circumstances for debugging errors concerning integration of doublet predictions from the ‘Remove’ and ‘Rescue’ steps.

#### Final Results

9. <u>“Final_doublets_groups_TEST.txt”</u> – This file contains the final list of predicted doublets and their associated new clusters following merging and regrouping. When using DoubletDecon as part of a processing pipeline with external applications this file will be of most use. The first column of the written file or the row names of the Main_Doublet_Decon() output are the final predicted doublets.
10. <u>“Final_nondoublets_groups_TEST.txt”</u> – This file is related to the above “Final_doublets_groups_TEST.txt” file, only with DoubletDecon’s final predicted non-doublets.
11. <u>“Final_doublets_exp_TEST.txt”</u> – This expression file is very similar to the “data_processed_TEST.txt” and the “data_processed_reclust_TEST.txt” files, but contains only DoubletDecon predicted doublets. It is important to note that the final expression files output by DoubletDecon are in ICGS/AltAnalyze compatible format and contain cluster identification rows and columns which may be incompatible with external pipelines. Removing these additional columns and rows in Excel or within R is recommended for reintegration with the Seurat pipeline.
12. <u>“Final_nondoublets_exp_TEST.txt”</u> – This expression file is related to the above “Final_doublets_exp_TEST.txt” file, only containing DoubletDecon’s final predicted non-doublets.

**Note**: Neither “data_processed_reclust” nor “groups_processed_reclust” are included in the function output of Main_Doublet_Decon(), only in the written results. This is because these files are unnecessary for downstream processing and do not provide new information for understanding the intermediate steps of DoubletDecon.

## LIMITATIONS

The performance and accuracy of DoubletDecon is dependent on the selection of key parameters of varying importance (listed in Step 3 and Step 4 of this protocol in descending order of importance). In addition to the dependence, DoubletDecon relies on a number of assumptions that may not hold true in all applications of the approach. Breaking of any of these assumptions may limit the applicability and performance of DoubletDecon. These include:

1. Representation of all cell states that contribute to doublet formation. For a specific doublet to be detected in DoubletDecon, a cell cluster for each of the two contributing cell types must be present in the dataset. Exclusion of a contributing cell type due to sorting, filtering, and other biological and computational means will limit DoubletDecon’s ability to predict doublets containing that cell type.
2. Accurate clustering of input data. DoubletDecon assumes that all clusters are transcriptionally distinct in that no two clusters have a highly similar transcriptional profile (i.e. subtypes of the same cell type) and that no cluster is truly a combination of two dissimilar clusters. Unsupervised clustering algorithms will frequently result in too granular or too coarse clustering, which must be mitigated by diligent checking of the cell clusters produced relative to what is biologically valid.
3. Homotypic doublets are relatively benign. Doublets occur when two cells of the same cell type are accidently sequenced together (homotypic doublets) and when two different cell types are erroneously combined (heterotypic doublets). DoubletDecon is only able to reliably detect heterotypic doublets, leaving datasets containing prevalent homotypic doublet populations sensitive to the effect of these doublets.
4. Mixed-lineage or transitional cell states will have unique gene expression. DoubletDecon works by identifying cells that are transcriptionally similar to doublets then rescuing erroneously identified cells, presumably transitional or mixed-lineage cells, based on expression of genes that are not highly expressed in the two contributing cell types. This relies on the assumption that mixed-lineage and transitional cells express genes that are related to the process of driving differentiation; however, this may not hold true in all circumstances.

## TROUBLESHOOTING

### Problem

You are receiving a “no locations are finite” error or output to the terminal stating “Unable to perform mcl function for blacklist clustering, please try a different rhop.” when running DoubletDecon.

### Potential Solution

This error most commonly occurs when the value you selected for the cluster merging parameter ρ’ (‘rhop’ in the parameter list) causes too much or too little merging of the clusters, leading to an error in the Markov clustering portion of the algorithm. Altering the value of this parameter according to the instructions given in Step 3 of this protocol should solve the issue, though incrementally stepping down from the ρ’ you are currently using (i.e. 1.1 to 1 to 0.9, etc.) will allow you to manually find the upper limit of acceptable ρ’ values. Alternatively, the DoubletDecon UI has a built-in function for displaying all possible values of ρ’ to make it easier to choose one within in the range. Once selected, make sure to evaluate the appropriateness of the clustering and adjust up or down depending on whether you want less or more cluster merging, respectively.

### Problem

You are receiving a “trying to get slot “var.genes” from an object of a basic class (“matrix”) with no slots” error from DoubletDecon when you are using Seurat input.

### Potential Solution

This error can occur because the RunPCA step in creating the Seurat object was run with all genes:

~~~
                 >pc.genes = rownames(object@data)
~~~

This causes an error during the synthetic doublet creation step of DoubletDecon. To potentially overcome this error, RunPCA in Seurat must be run using variable genes:

~~~
                 >pc.genes = object@var.genes
~~~

### Problem

You are receiving an “Error in socketConnection(“localhost”, port = port, server = TRUE, blocking = TRUE,: all connections are in use” error when running DoubletDecon.

### Potential Solution

This error can occur when using the automatic detection option for the number of cores available in the rescue step of DoubletDecon (‘nCores’ = 1). While this error can arise due to specific settings on a personal computer, it is more likely to affect the DoubletDecon run when using a computational cluster with more cores technically available than is assigned to the specific user. To overcome this, turn off automatic detection and manually set the number of cores using the ‘nCores’ parameter.

## ACKNOWLEDGMENTS

This work was partly funded by CCHMC Research Foundation (NS).

## AUTHOR CONTRIBUTIONS

Writing, E.A.K.D, D.S., K.C., N.S.; Development, E.A.K.D, D.S, and N.S.; Processing, E.A.K.D., D.S., K.C.; Funding Acquisition, N.S.

## KEY RESOURCES TABLE

| REAGENT or RESOURCE | SOURCE | IDENTIFIER |
| --- | --- | --- |
| Deposited Data |  |  |
| Single-cell RNA-sequencing data (bone marrow progenitors) | GEO | GEO: GSE70245 |
| Single-cell RNA-sequencing data (PBMCs demultiplexed using Demuxlet) | GEO | GEO: GSE96583 |
| Software and Algorithms |  |  |
| DoubletDecon | DePasquale et al., 2019 | https://github.com/EDePasquale/DoubletDecon |
| AltAnalyze (ICGS) | Olsson et al., 2016 | http://altanalyze.org |
| Seurat | Satija et al., 2015 | https://satijalab.org/seurat |

